# Chemogenetic disconnection between the orbitofrontal cortex and the rostromedial caudate nucleus disrupts motivational control of goal-directed action

**DOI:** 10.1101/2022.04.22.489147

**Authors:** Kei Oyama, Yukiko Hori, Koki Mimura, Yuji Nagai, Mark A G Eldridge, Richard C Saunders, Naohisa Miyakawa, Toshiyuki Hirabayashi, Yuki Hori, Ken-ichi Inoue, Tetsuya Suhara, Masahiko Takada, Makoto Higuchi, Barry J Richmond, Takafumi Minamimoto

## Abstract

The orbitofrontal cortex (OFC) and its major downstream target within the basal ganglia—the rostromedial caudate nucleus (rmCD)—are involved in reward-value processing and goal-directed behavior. However, a causal contribution of the pathway linking these two structures to goal-directed behavior has not been established. Using the chemogenetic technology of Designer Receptors Exclusively Activated by Designer Drugs with a crossed inactivation design, we functionally and reversibly disrupted interactions between the OFC and rmCD in two male macaque monkeys. We injected an adeno-associated virus vector expressing an inhibitory designer receptor (hM4Di) into the OFC and contralateral rmCD, the expression of which was visualized *in vivo* by positron emission tomography (PET) and confirmed by post-mortem immunohistochemistry. Functional disconnection of the OFC and rmCD resulted in a significant and reproducible loss of sensitivity to the cued reward value for goal-directed action. This decreased sensitivity was most prominent when monkeys had accumulated a certain amount of reward. These results provide causal evidence that the interaction between the OFC and the rmCD is needed for motivational control of action on the basis of the relative reward value and internal drive. This finding extends current understanding of the physiological basis of psychiatric disorders in which goal-directed behavior is affected, such as obsessive-compulsive disorder.

**Significance Statement:** In daily life, we routinely adjust the speed and accuracy of our actions on the basis of the value of expected reward. Abnormalities in these kinds of motivational adjustments might be related to behaviors seen in psychiatric disorders such as obsessive-compulsive disorder. In the current study, we show that the connection from the orbitofrontal cortex to the rostromedial caudate nucleus is essential formotivational control of action in monkeys. This finding expands our knowledge about how the primate brain controls motivation and behavior and provides a particular insight into disorders like obsessive-compulsive disorder, in which altered connectivity between the orbitofrontal cortex and the striatum has been implicated.

## Introduction

The subjective desirability (i.e., value) of an expected reward is a key factor in determining the latency, accuracy, and vigor of goal-directed behavior (Dickinson and Balleine, 1994). Goal-directed behavior is regulated by two factors: the incentive value of the goal (reward) and the internal drive (physiological state) of an agent (Berridge, 2004; Zhang et al., 2009). The orbitofrontal cortex (OFC) is thought to be critical for the animal’s ability to adjust behavior based on reward value. Neuronal activity in the OFC is substantially modulated by the sensory and hedonic properties of rewards (Rolls et al., 1989; de Araujo and Rolls, 2004) as well as subjective reward preference (Padoa-Schioppa and Assad, 2006; Chaudhry et al., 2009). Inactivation or lesion of the bilateral OFC disrupts the ability to use stimulus information for directing behaviors to optimize the outcome (Izquierdo et al., 2004; Murray et al., 2015). However, it remains unclear which downstream region receives the value-related information from the OFC and processes it for implementing goal-directed behavior.

Although the ventral part of the striatum is generally regarded as the primary basal ganglia destination downstream of the OFC, anatomical studies in monkeys have reported that the rostromedial part of the caudate nucleus (rmCD) also receives direct ipsilateral connections from the OFC, specifically Brodmann areas (BA) 11 and 13 (Haber et al., 2006). Electrophysiological recording studies in monkeys have reported that neuronal activity in the rmCD signals information about the expected reward value and satiation level, but exhibits a relatively weak selectivity for movements (Hollerman et al., 1998; Nakamura et al., 2012; Fujimoto et al., 2019). Additionally, temporary inactivation of bilateral—but not unilateral— rmCD neuron activity impaired the ability of monkeys to adjust their motivation based on incentive cues (Nagai et al., 2016). Moreover, research in rodents has demonstrated that the projection from the OFC to the dorsomedial striatum is a critical pathway for carrying the incentive information and behaving in a goal-directed manner (Yin et al., 2005; Gremel and Costa, 2013; Gremel et al., 2016). These findings thus provide a plausible scenario in which the primate OFC-rmCD projection contributes to the motivational adjustment of action based on incentive and drive; however, this has not yet been directly examined. Addressing this issue is particularly important considering that disrupted functional connectivity between the OFC and the striatum is implicated in many human psychiatric disorders, including obsessive compulsive disorder (OCD), whose symptoms seem to be associated with impaired motivational control of behavior (Harrison et al., 2009; Figee et al., 2013; Abe et al., 2015; Jahanshahi et al., 2015; Gillan et al., 2016).

To determine the contribution of the OFC-rmCD pathway in goal-directed behavior, here we used a chemogenetic technology—Designer Receptors Exclusively Activated by Designer Drugs (DREADDs)—with a crossed inactivation design to reversibly disrupt their direct intrahemispheric information flow. We virally introduced an inhibitory designer receptor (hM4Di) into OFC and contralateral rmCD neurons in two macaque monkeys. We used a reward-size task to examine the effect of temporarily disconnecting these areas on monkeys’ ability to adjust goal-directed actions based on motivation. The relationships between motivational value, incentive, and drive were inferred from task performance. We show that following systemic administration of DREADD agonists, goal-directed behavior in monkeys was altered such that they became insensitive to differences in reward magnitude and satiation.

## Materials and Methods

### Animals

Two male macaque monkeys participated in the experiments (MK#1: Rhesus monkey [*Macaca mulatta*] 7.3 kg, aged 10.3 years at the start of experiments; MK#2: Japanese monkey [*Macaca fuscata*] 6.1 kg, aged 4.4 years at the start of experiments). All experimental procedures involving animals were carried out in accordance with the Guide for the Care and Use of Nonhuman primates in Neuroscience Research (The Japan Neuroscience Society; https://www.jnss.org/en/animal_primates) and were approved by the Animal Ethics Committee of the National Institutes for Quantum Science and Technology. A standard diet, supplementary fruits/vegetables, and a tablet of vitamin C (200 mg) were provided daily.

### Viral vector production

AAV2 vectors (AAV2-CMV-hM4Di and AAV2-CMV-AcGFP) were produced by helper-free triple transfection and purified by affinity chromatography (GE Healthcare, Chicago, IL, USA). Viral titer was determined by quantitative polymerase chain reaction using Taq-Man technology (Life Technologies, Waltham, MA, USA).

### Surgical procedures and viral vector injections

Surgeries were performed under aseptic conditions in a fully equipped operating suite. We monitored body temperature, heart rate, SpO_2_, and tidal CO_2_ throughout all surgical procedures. Anesthesia was induced using an intramuscular (i.m.) injection ofketamine (5–10 mg/kg) and xylazine (0.2–0.5 mg/kg) and monkeys were intubated with an endotracheal tube. Anesthesia was maintained with isoflurane (1%–3%, to effect). After surgery, prophylactic antibiotics and analgesics were administered. Before surgery, magnetic resonance (MR) imaging (7 Tesla 400 mm/SS system, NIRS/KOBELCO/Brucker) and X-ray computed tomography (CT) scans (Accuitomo170, J. MORITA CO., Kyoto, Japan) were acquired under anesthesia (continuous infusion of propofol 0.2–0.6 mg/kg/min, intravenously). Overlaid MR and CT images were created using PMOD® image analysis software (PMOD Technologies Ltd., Zurich, Switzerland) to estimate stereotaxic coordinates of target brain structures.

The monkeys were first co-injected with AAV2-CMV-hM4Di (1.0 × 10^13^ and 2.3 × 10^13^ particles/mL,respectively)and AAV2-CMV-AcGFP (4.7 × 10^12^ and 6.6 × 10^12^ particles/mL, respectively) into the OFC of one hemisphere (BA11 & BA13; right and left hemispheres, respectively) (Fig. 1B, C). The injections were performed under direct vision. The OFC was visualized using the same types of surgical procedures as used as a previous study (Eldridge et al., 2016). Briefly, after retracting skin, galea, and muscle, the frontal cortex was exposed by removing a bone flap and reflecting the dura mater. Handheld injections were then made under visual guidance through an operating microscope (Leica M220, Leica Microsystems GmbH, Wetzlar, Germany), with care taken to place the beveled tip of a microsyringe (Model 1701RN, Hamilton) containing the viral vector at an angle oblique to the brain surface. The needle (26 Gauge, PT2) was inserted into the intended area of injection by one experimenter and a second experimenter pressed the plunger to expel approximately 1 μL per penetration. Totals of 54 μL and 50 μL were injected for MK#1 and MK#2 via 53 and 49 tracks, respectively.

**Figure 1.**
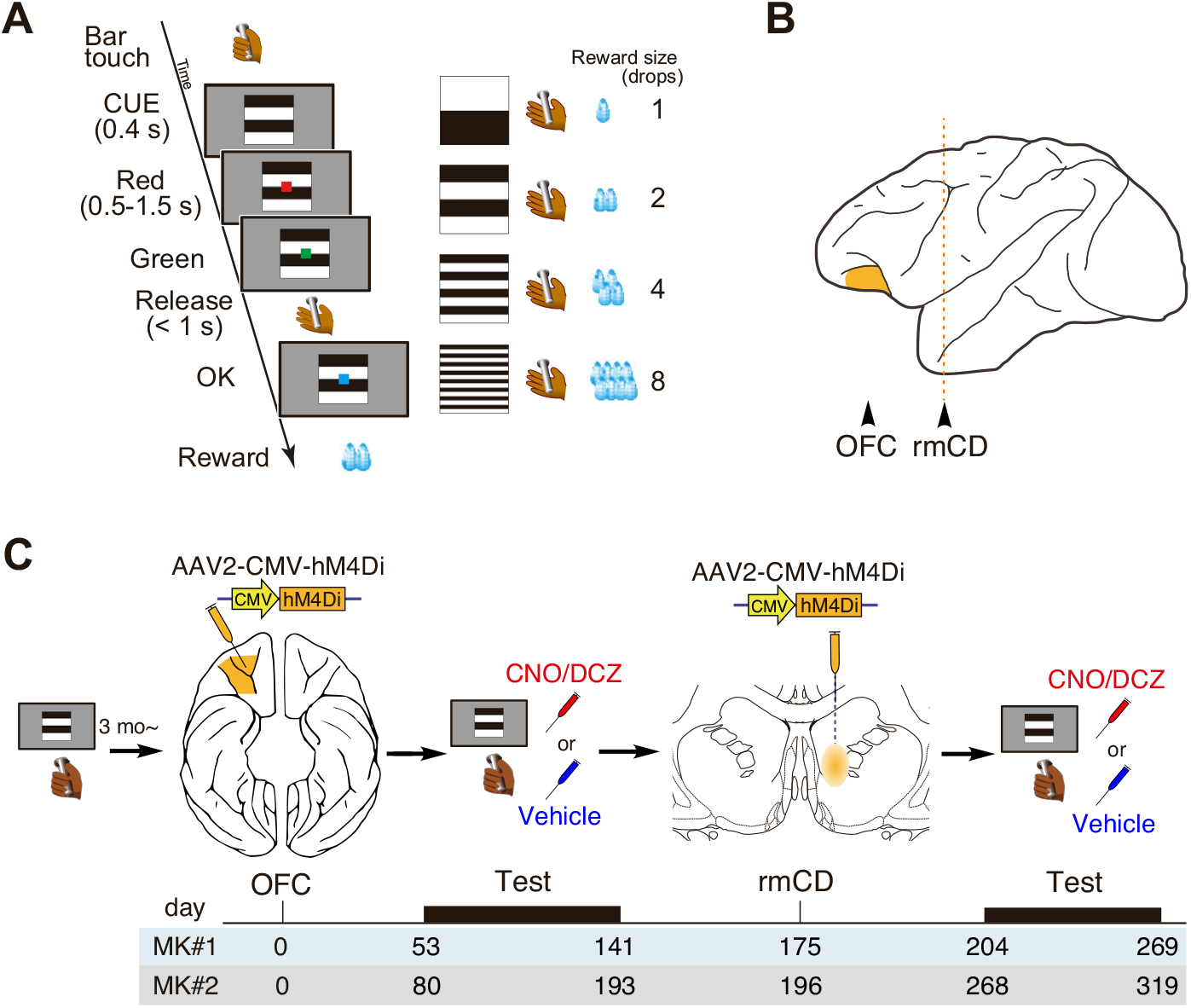
Task and experimental design. (A) The reward-size task. Left: At the beginning of each trial, a visual cue signaled the amount of reward (1, 2, 4, or 8 drops) that would be delivered after a correct red/green color discrimination. After each correct trial, a new cue-reward size pair was picked from the set of four at random. Right: Relationship between cues and reward sizes. (B) Lateral view of monkey brain. Locations ofthe OFC and rmCD are indicated by arrow heads. (C) Experimental design and timeline. The monkeys were first trained on the reward-size task, followed by injections of the hM4Di vector into the OFC (purple) to produce unilateral OFC silencing when DREADD agonists were administered. Another AAV vector was then injected into the contralateral rmCD to enable functional disconnection of the OFC-rmCD pathway when the DREADD agonists were administered. The numbers at the bottom indicate the days after OFC injection.

At 174 and 196 days after the first injection, the monkeys received a second set of injections with the same vectors into the rmCD contralateral to the OFC injections (Fig. 1B, C) using a procedure used in a previous study (Nagai et al., 2016). Briefly, viruses (total volume, 3 μL for both monkeys) were pressure-injected by a 10-μL microsyringe (Model 1701RN, Hamilton) with a 30-gauge injection needle in a fused silica capillary (450 mm OD) to create a step approximately 500 μm away from the needle tip to minimize backflow. The microsyringe was mounted into a motorized microinjector (UMP3T-2, WPI) that was held by a manipulator (Model 1460, David Kopf, Ltd.) on the stereotaxic frame. After a burr hole (8-mm diameter) and the dura mater (~5 mm) were opened, the injection needle was inserted into the brain and slowly moved down 2–3 mm beyond the target, then kept stationary for 5 min, after which it was pulled up to the target location. The injection speed was set at 0.5 μL/min. After the injection, the needle remained *in situ* for 15 min to minimize backflow along the needle.

### Behavioral task

Before the experiments, monkeys were trained for more than 3 months to discriminate the color of shapes presented on a computer monitor. For the experiment, we used a reward-size task, as described previously (Minamimoto et al., 2009) (Fig. 1A), which followed a cued multi-trial reward schedule. Task control and data acquisition were performed using a QNX-based real-time experimentation data acquisition system (REX; Laboratory of Sensorimotor Research, NEI/NIH) and commercially available software (Presentation, Neurobehavioral Systems). The use of an automated system eliminated the need for experimenters to be blind to treatment. A monkey initiated a trial by touching a bar. A background visual cue and a red target appeared sequentially on the display. After a variable interval, the red target turned green. If the monkey released the bar between 200 and 1,000 ms after this event, the target turned blue and a liquid reward (1, 2, 4, or 8 drops; 1 drop = ~0.1 mL) was delivered immediately afterward. If the monkey released the bar outside the 200-1,000 ms range, we regarded the trial as an “error” trial, which was aborted and repeated after a 1,000-ms inter-trial interval. The visual cue presented at the beginning of the trial indicated the amount of reward that would be received if the trial was completed successfully. After an error, the monkey had to repeat the same trial condition and correctly complete it to receive a reward. Because the monkeys were able to perform the task correctly on nearly every trial when the visual cues did not convey the information about the amount of reward, error trials were interpreted as those in which the monkeys were not sufficiently motivated to release the bar correctly (Minamimoto et al., 2009). Our behavioral measure for the expected outcome value was the proportion of error trials (i.e., error rates). Before each 100-min testing session, the monkeys went without water for approximately 22 h. Both monkeys were trained on the reward-size task for at least 3 weeks before the experiments.

### Drug administration

We used clozapine N-oxide (CNO) and deschloroclozapine (DCZ) as DREADD actuators for MK#1 and MK#2, respectively. CNO (Toronto Research) and DCZ (MedChemExpress) were dissolved in 2.0% or 2.5% dimethyl sulfoxide (DMSO, FUJIFILM Wako Pure Chemical Co.). These stock solutions were diluted in saline to a final volume of 2 mL at a dose of 3 mg/kg (CNO) or 1 mL at a dose of 0.03 mg/kg (DCZ) and injected intravenously (i.v.) or i.m. 15 min before the beginning of the experiments, respectively. These doses of CNO or DCZ yield 50%–60% hM4Di occupancy but do not affect the performance of monkeys which are not expressing hM4Di (Nagai et al., 2016, 2020). CNO/DCZ and vehicle treatment were tested no more than once per week.

### Positron emission tomography (PET) imaging

To examine the expression of hM4Di *in vivo*, positron emission tomography (PET) imaging was conducted before vector injection and approximately 8 weeks after injection for MK#2, as previously reported (Nagai et al., 2016, 2020). Briefly, the monkey was anesthetized with ketamine (5–10 mg/kg, i.m.) and xylazine (0.2–0.5 mg/kg, i.m.), which was maintained with isoflurane (1%—3%) during all PET procedures. PET scans were performed using a microPET Focus 220 scanner (Siemens Medical Solutions, Malvern, PA, USA). Transmission scans were performed for approximately 20 min with a ^68^Ge source. Emission scans were acquired in 3D list mode with an energy window of 350–750 keV after an intravenous bolus injection of [“C]DCZ (324.5–384.9 MBq). Emission-data acquisition lasted 90 min. PET image reconstruction was performed with filtered back-projection using a Hanning filter cut-off at a Nyquist frequency of 0.5 mm^−1^. To estimate the specific binding of [^11^C]DCZ, the regional binding potential relative to non-displaceable radioligand (BP_ND_) was calculated using PMOD® with an original multilinear reference tissue model (MRTMo) and the cerebellum as a reference (Nagai et al., 2016; Yan et al., 2021). Contrast (subtraction) images were constructed using SPM12 software (Welcome Department of Imaging Science; www.fil.ion.ucl.ac.uk) and MATLAB R2016a software (MathWorks Inc., Natick, MA, USA). The surface of the gray matter was estimated on the basis of the MR image. Briefly, Yerkes standard T1 and T2 templates (Donahue et al., 2016, 2018) were first linearly and nonlinearly registered to the original MR image using FMRIB’s linear registration tool (FLIRT) and FMRIB’s nonlinear registration tool (FNIRT) in FSL software (Smith et al., 2004). The gray matter surface was then estimated using registered templates and the HCP-NHP structural pipeline (Autio et al., 2020). Finally, a contrast PET image was projected on the gray matter surface using Connectome Workbench software (https://www.humanconnectome.org/software/get-connectome-workbench), followed by registration of the contrast PET image to the original MR image.

### Histology and immunostaining

For histological inspection, two monkeys were deeply anesthetized with an overdose of sodium pentobarbital (80 mg/kg, i.v.) and transcardially perfused with saline at 4 °C, followed by 4% paraformaldehyde in 0.1 M phosphate buffered saline (PBS) with a pH of 7.4. The brain was removed from the skull, postfixed in the same fresh fixative overnight, saturated with 30% sucrose in phosphate buffer (PB) at 4°C, then cut serially into 50-μm-thick sections on a freezing microtome. For visualization of immunoreactive signals of GFP co-expressed with hM4Di, every 6th section was immersed in 1% skim milk for 1 h at room temperature and incubated overnight at 4 °C with rabbit anti-GFP monoclonal antibody (1:200–500, G10362, Thermo Fisher Scientific), then for 2 days in PBS containing 0.1% Triton X-100 and 1% normal goat serum at 4 °C. The sections were then incubated in the same fresh medium containing biotinylated goat anti-rabbit IgG antibody (1:1,000; Jackson Immuno Research, West Grove, PA, USA) for 2 h at room temperature, followed by avidin-biotin-peroxidase complex (ABC Elite, Vector Laboratories, Burlingame, CA, USA) for 2 h at room temperature. For visualization of the antigen, the sections were reacted in 0.05 M Tris-HCl buffer (pH 7.6) containing 0.04% diaminobenzidine (DAB), 0.04%NiCl_2_, and 0.003% H_2_O_2_. The sections were then mounted on gelatin-coated glass slides, air-dried, and cover-slipped. A second series of sections was Nissl-stained with 1% Cresyl violet. Images of sections were digitally captured using an optical microscope equipped with a high-grade charge-coupled device camera (BZ-X710, Keyence, Osaka, Japan) or a whole slide scanner (NanoZoomer S60, Hamamatsu Photonics, Hamamatsu, Japan).

### Experimental Design and Statistical Analysis

For behavioral data analysis, the error rate for each reward size was calculated for each daily session. We used the error rates to estimate the level of motivation, as the error rates of these tasks (*E*) were inversely related to the value for action. In the reward-size task, we used an inverse function,

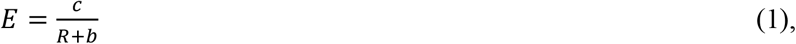

where *E* is error rate, *R* is reward size, and *c* and *b* are constants.

To examine the effects of satiation, we divided each session into quartiles based on normalized cumulative reward, *R_cum_*, which was 0.125, 0.375, 0.625, and 0.875 for the first through fourth quartiles, respectively. We fitted the error rates obtained from each monkey to the following model:

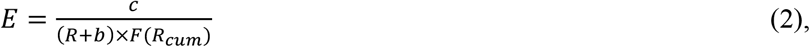

where the satiation effect, *F(R_cum_*), represents the exponentially decaying reward value (at a constant rate) as reward accumulates (i.e., *R_cum_* increases) (Minamimoto et al., 2012):

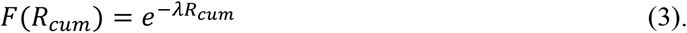

We fitted the functions to the data using sum-of-squares minimization. We performed repeated-measures analysis of variance (ANOVA) with individual monkeys nested to test the effect of treatment and its interactive effect with reward size and satiation on error rates. For example, the effect of treatment × reward size on error rates was examined using an R code: aov(error ~ treatment * reward + Error[subject], data). Separate ANOVAs were also conducted on each subject’s data for confirmation.

## Results

### Unilateral silencing ofthe OFC had little effect on reward-size task performance

Both monkeys were injected with AAV vectors expressing hM4Di in unilateral OFC (BA11 & BA13, Fig. 1C). Eight weeks after the injection, we visualized the expression of the DREADD *in -vivo* via PET imaging using the DREADD-selective radioligand ^11^C-labcled DCZ (Nagai et al., 2020). As shown in Fig. 2AB, an increase in PET signal covered the target region of the OFC, which indicated hM4Di expression. Expression in individual OFC neurons was verified using post-mortem immunohistochemical staining for the co-expressed AcGFP (e.g., Fig. 2C), at an anterior-posterior level corresponding to Fig. 2AB−1 to −3 in both monkeys.

**Figure 2.**
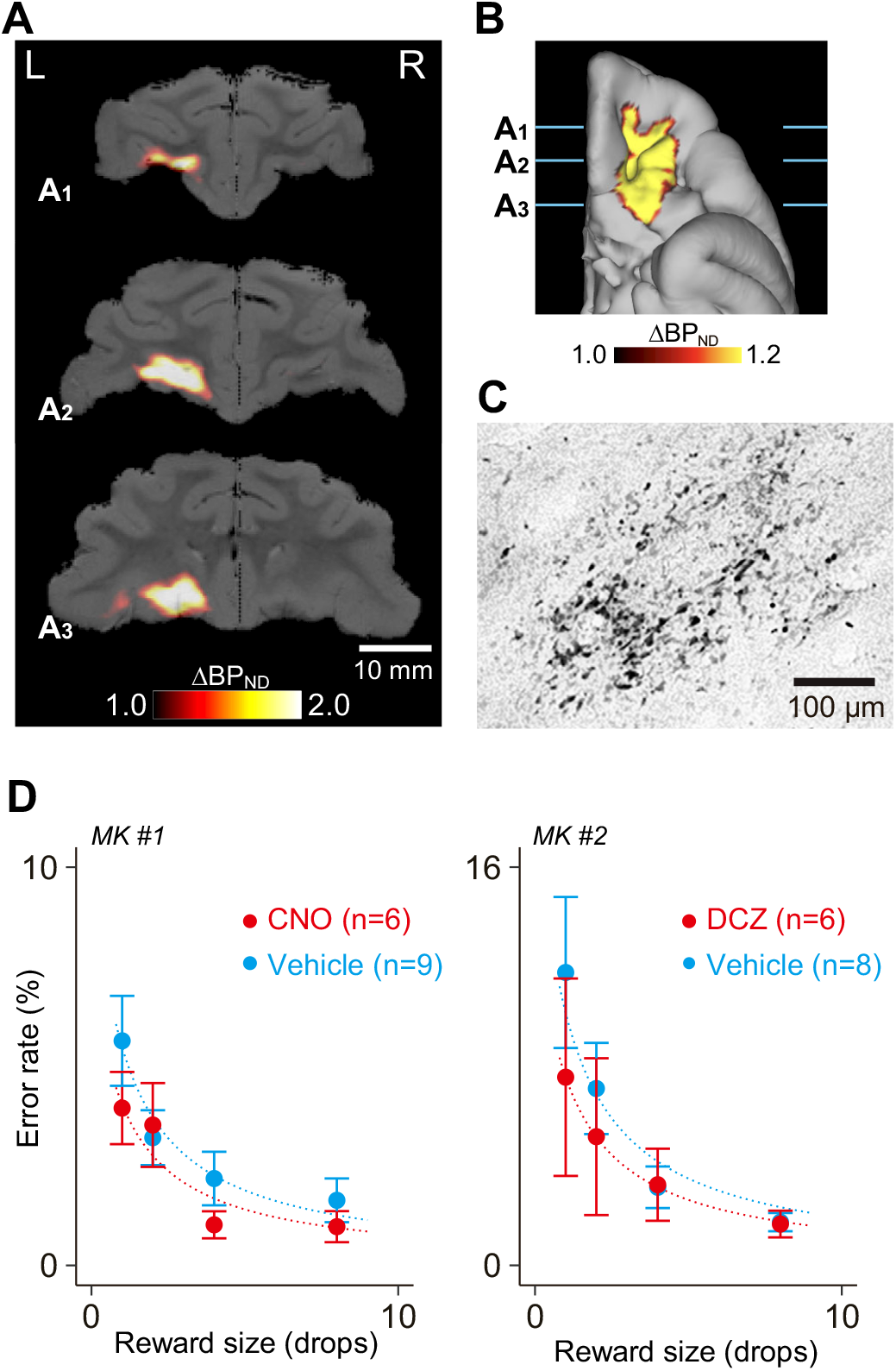
Effect of unilateral chemogenetic OFC inactivation on reward evaluation. (AB) *In vivo* visualization ofhM4Di in the OFC using PET. Coronal parametric image (Al–A3, anterior to posterior) and ventral view of flat map (B) showing increased tracer [^11^C]DCZ binding (BP_ND_) 58 days after hM4Di vector injection compared with pre-vector injection, overlaid on the MR image for MK#2. (C) Bright-field image of a coronal DAB-stained section showing GFP-positive neurons in the OFC. (D) Effect of chemogenetic silencing of unilateral OFC on performance during the reward-size task in MK#1 (left) and MK#2 (right), respectively. Error rates as a function of reward size (mean ± SEM) are plotted for DREADD agonist (red) and vehicle (cyan) treatment conditions. Curves are the best fit function of Eq. 1. We found no significant effect of silencing on error rates in either monkey (MK#1, F_(1, 52)_ = 1.9, *p* = 0.17; MK2, F_(1,48)_ = 0.98, *p* = 0.33).

To examine the ability to estimate reward and adjust behavior, we tested the monkeys on the reward-size task (Fig. 1A). The task requirement (i.e., release the bar on time) was so simple that after 3 months of training, the monkeys could complete high incentive trials with a near-100% rate of accuracy if sufficiently motivated (Minamimoto et al., 2009). Errors (either releasing the bar too early or too late) were typically observed in small reward trials and/or close to the end of daily sessions and were therefore considered as a behavioral indicator that the monkeys were not sufficiently motivated to correctly release the bar. As shown in earlier studies (e.g., Minamimoto et al., 2009), the error rates were related to the value of the upcoming reward (Fig. 2D), with fewer errors occurring when expected reward was high. Overall error rates after treatment with the DREADD agonists CNO or DCZ did not differ from those after injection with vehicle control (two-way ANOVA, main effect of treatment, F_(1, 107)_ = 2.1, *p* = 0.16; Fig. 2D). We consistently observed a significant effect of reward on error rates (main effect of reward size, F_(3, 107)_ = 10.9, *p* = 2.7 × 10^−6^), but not of the interaction (reward × treatment, F_(3, 107)_ = 0.49, *p* = 0.69). This result suggests that unilateral silencing of the OFC did not interfere with the normal ability to estimate reward, which is in accordance with a previous study of OFC lesions (Clark et al., 2013).

Unilateral OFC inactivation did not alter the frequency of error types (early or late errors) (two-way ANOVA, main effect of treatment, F_(1, 26)_ = l.5, *p* = 0.24). The treatment significantly shorten reaction time (two-way ANOVA; treatment: F_(1, 107)_ = 43.1, *p* = 1.9 × 10^−9^). Reward size also had significant impact on reaction time (F_(3, 107)_ = 9.2, *p* = 1.9 × 10^−5^), but without significant interaction with treatment (reward × treatment: F_(3,107)_ = 0.08, *p* = 0.97). Total reward earned significantly increased following inactivation (F_(1, 26)_ = 6.0, *p* = 0.02). However, no significant interaction was observed between treatment and satiation on error rates (three-way ANOVA with treatment, reward size, and satiation; treatment × satiation, F_(3, 31)_ = 0.81, *p* = 0.50)(see Discussion).

### Chemogenetic disconnection of OFC-rmCD reduced the sensitivity to the amount ofexpected reward

Next, we examined the causative role of the functional connection between the OFC and the rmCD by contralateral (crossed) inactivation of these two areas. After OFC vector injection, PET showed increased binding of [^11^C]DCZ in the ipsilateral rmCD (Fig. 3A, left), reflecting hM4Di-positive axon terminals (i.e., anatomical connection from the OFC to the rmCD as reported in previous anatomical studies; Haber et al., 2006). Increased binding of [^11^C]DCZ was also found in the ipsilateral medial part of the mediodorsal thalamus (Fig. 3B). We then injected the viral vector into the rmCD contralateral to the first injections (Fig. 1C). The [^11^C]DCZ-PET scans detected the expression of hM4Di in the rmCD as increased tracer binding extending 2–3 mm anterior to posterior, resulting in a mirror image of the OFC terminal site (Fig. 3A). This was further verified by post-mortem immunohistochemical staining for the co-expressed AcGFP for both monkeys (e.g., Fig. 3C).

**Figure 3.**
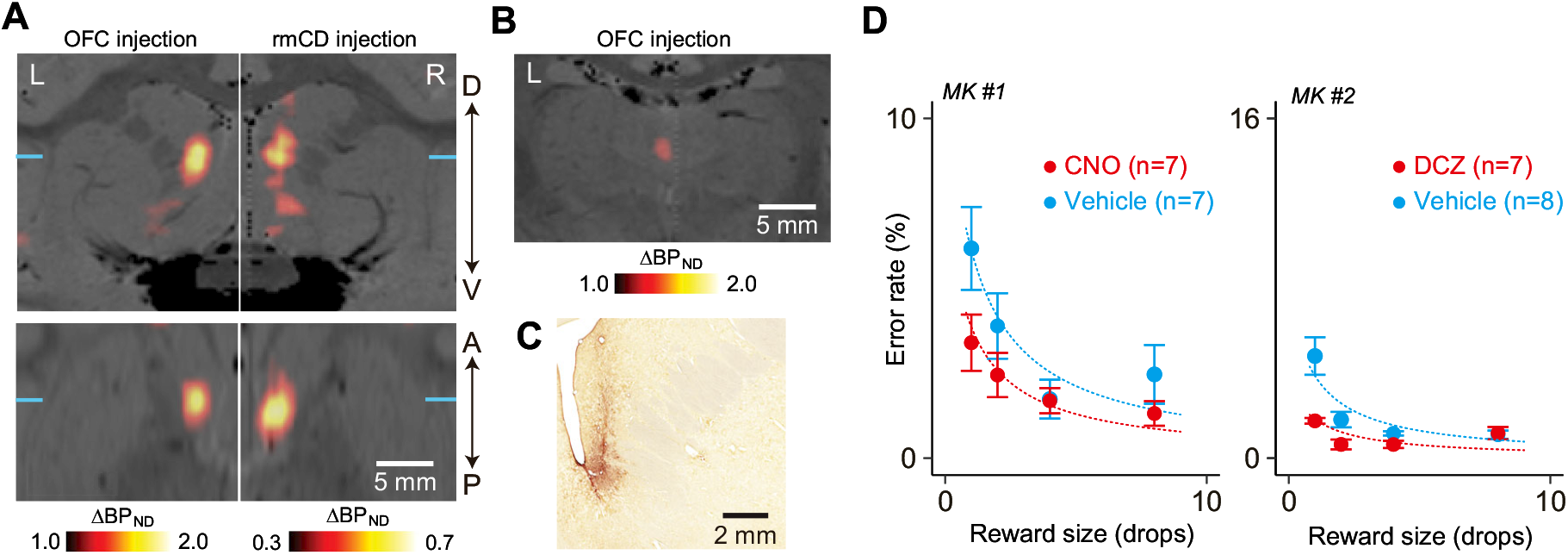
Chemogenetic disconnection of OFC-rmCD reduced the sensitivity to reward amount. (A) *In vivo* visualization of hM4Di in the rmCD using PET. Coronal (top) and horizontal PET images (bottom) representing increased relative tracer binding (BP_ND_) 58 days after vector injection into the OFC (OFC injection, left) and 57 days after injection into the rmCD (rmCD injection, right) compared with pre-vector injection, overlaid on an MR image for MK#2. Each coronal and horizontal section was acquired at the anterior-posterior (A-P) and dorsal-ventral (D-V) levels indicated by corresponding blue lines, respectively. (B) Coronal PET image representing increased BP_ND_ in the mediodorsal thalamus, indicating hM4Di-positive OFC axon terminals. (C) Bright-field image of a coronal DAB-stained section showing immunoreactivity against a reporter protein (GFP) in the rmCD for MK#2. (D) Effect of chemogenetic functional disconnection of the OFC and rmCD on performance during the reward-size task in MK#1 (left) and MK#2 (right), respectively. Error rates as a function of reward size (mean ± SEM) are plotted for DREADD agonist (red) and vehicle (cyan) treatment conditions. Curves are the best fit function of Eq. 1. A significant main effect of treatment was observed in both monkeys (MK#l, F_(1,48)_ = 6.0, *p* = 0.018; MK#2, F_(1,52)_ = 17.1, *p* = 1.3×10^−4^).

We then tested the monkeys on the reward-size task following treatment with the DREADD agonists or vehicle control. We found that error rates were consistently lower after treatment with the DREADD agonists compared with those after treatment with vehicle controls (Fig. 3D; two-way ANOVA, main effect of treatment, F_(1,107)_ = 17.7, *p* = 5.4 × 10^−5^). The treatment had a significant interaction effect with reward size (reward × treatment, F_(3,107)_ = 3.9, *p* = 0.011)— such that treatment reduced error rates more for trials in which expected rewards should have been small, while the impact of reward on error rates remained significant (main effect of reward size, F_(3,107)_ =17.7, *p* = 2.1 × 10^−9^). Considering that the unilateral inactivation of either OFC (Fig. 2D) or rmCD alone (Nagai et al., 2016) did not change error rates on this task, our results suggest that functional disconnection between OFC and rmCD reduced the sensitivity to differences in reward size.

We further examined the effect of the OFC-rmCD chemogenetic disconnection on other behavioral parameters. We found that disconnection significantly increased the early/late error ratio (main effect of treatment, F_(1, 26)_ = 11.2, *p* = 0.0025). The disconnection tended to shorten reaction time (two-way ANOVA; main effect of treatment, F_(1,107)_ = 3.9, *p* = 0.051), while the impact of reward remained significant (main effect of reward size, F_(3,107)_ = 3.8, *p* = 0.012). The treatment had no significant interactive effect with reward size on reaction time (treatment × reward size, F_(3,107)_ = 0.6, *p* = 0.60).

### Chemogenetic disconnection of OFC-rmCD reduced the impact ofsatiation on performance

The motivational value of reward should decrease as the physiological drive state changes from thirst to satiation. Because it has been shown that this devaluation effect on goal-directed action is diminished after OFC lesions (e.g., Izquierdo et al., 2004), the decreased sensitivity to reward magnitude that we observed might have resulted from decreased sensitivity to satiation shift. In all daily sessions, the monkeys were allowed to keep performing the task until they did not want to anymore, meaning that the final data each day were collected as the monkeys were approaching satiation. When treated with the vehicle, overall error rates for each reward size increased as the normalized cumulative reward (*R_cum_*) increased (Fig. 4, left). In contrast, satiation-dependent changes in error rates were not pronounced following treatment with the DREADD agonists (Fig. 4, right). Indeed, we observed a significant interaction between treatment and satiation on error rates (three-way ANOVA with treatment, reward size, and satiation; treatment × satiation, F_(3,31)_ = 9.06, *p* = 1.9× 10^−4^). As OFC-rmCD disconnection did not change the amount of total reward received (F_(1,26)_ = 2.l8, *p* = 0.15), the drive for wateror general motivation to perform the task did not seem to have been altered. Collectively, these data suggest that OFC-rmCD disconnection significantly attenuated the impact of satiation on goal-directed performance.

**Figure 4.**
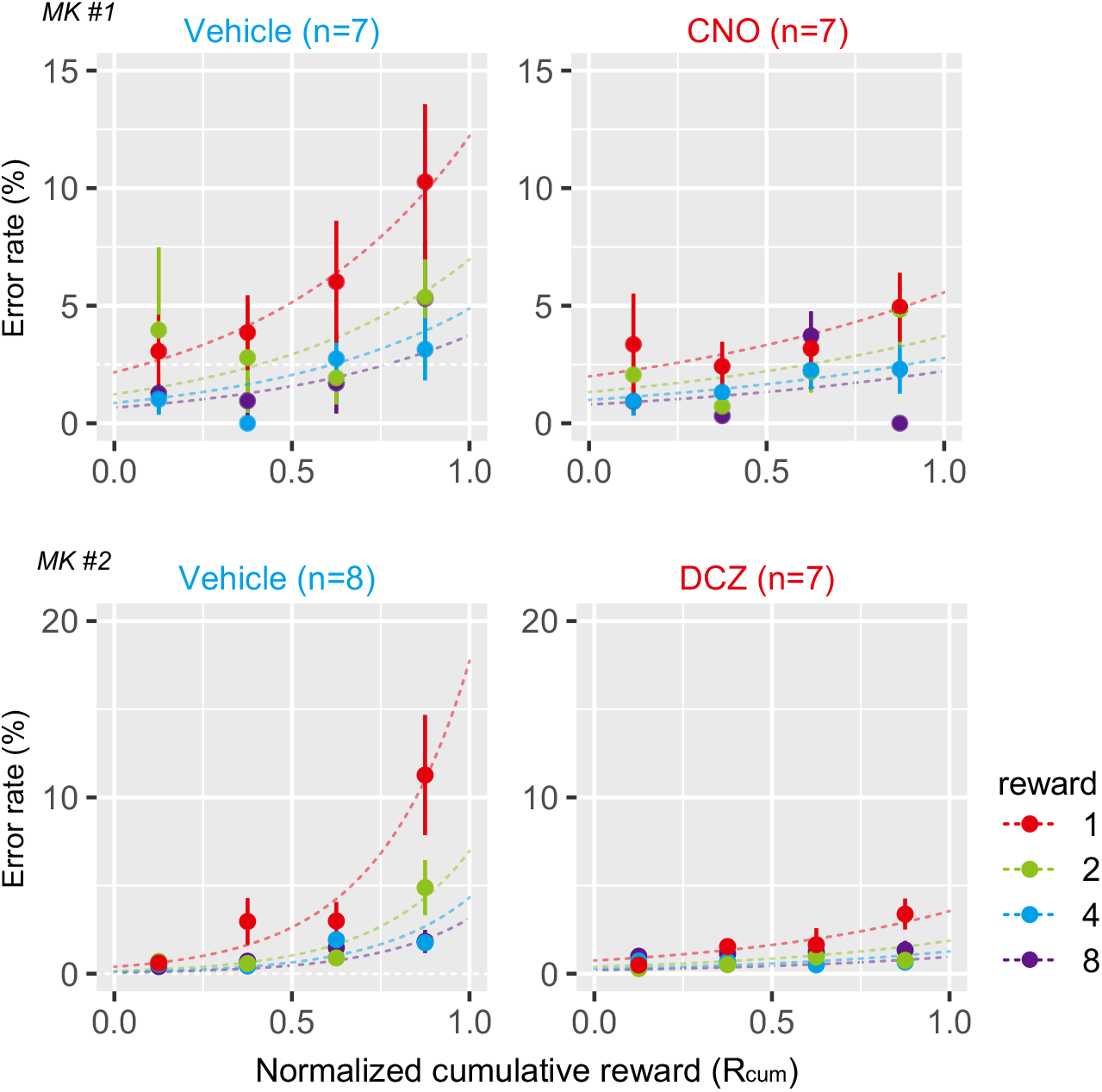
Chemogenetic disconnection of OFC-rmCD reduced the impact of satiation on performance. Error rates as a function of normalized cumulative reward (mean ± SEM) are plotted for vehicle (left) and DREADD agonist treatment conditions (right) in MK#l (top) and MK#2 (bottom), respectively. Reward condition (amount) is color coded. Curves are the best fit function of Eq. 2.

## Discussion

To examine the causal role of the communication through the OFC-rmCD pathway in goal-directed behavior, we used DREADD technology to functionally and temporarily disconnect these brain areas in two macaques. Activation ofhM4Di in the OFC and the contralateral rmCD produced a significant and reproducible loss of normal sensitivity to the cued reward value for goal-directed action. The disconnection did not decrease general attention or drive, as evidenced by the unaffected reaction times and the total amount of reward earned. Reduced sensitivity to reward size was most prominent when monkeys had accumulated a certain amount of reward, suggesting that the satiation effect on motivation depends on the integrity of the OFC-rmCD connections.

Expectation of the reward value is the hallmark of motivational control of behavior. The motivational value comprises both external incentives and internal drives. In the reward-size task, visual cues always indicated the amount of reward for a given trial. At the same time, the motivation to receive the reward—indexed by bar-releasing error rates—became less as the monkeys drank throughout the session. Thus, the motivational value of the cue was dynamically updated according to the degree of satiation. Previous studies using the reward-size task have shown that the OFC and rmCD control normal estimates of the cued outcome value. Specifically, monkeys with OFC ablation performed this task with smaller differences in error rates, suggesting a less sensitivity to relative reward amount (Simmons et al., 2010). When the rmCD was inactivated bilaterally, by using DREADDs or muscimol, the overall error rates increased and their discrimination of reward sizes diminished, indicating the decreased sensitivity to absolute and relative value estimation, respectively (Nagai et al., 2016). In these cases, however, the impact of satiation on performance remained normal. In contrast to these previous studies, the present work demonstrated that the OFC-rmCD disconnection reduced the impact of reward magnitude and satiation, and resulted in seemingly higher motivation when the reward value became small. In the same monkeys, however, the impact of incentive or satiation was unchanged when the unilateral OFC alone was silenced. Although the current study did not compare the behavioral effects of two silencing conditions directly, our results, together with the non-significant effects of unilateral rmCD silencing (Nagai et al., 2016), support the notion that the effects of OFC-rmCD disconnection is not the sum of two unilateral effects. Thus, our findings extend those of previous reports in an important way, indicating that communication from the OFC to the rmCD is critical for value updating and/or adjusting behavior based on internal drive.

Neurons in the OFC are known to represent the values of reward-predicting stimuli (Roesch and Olson, 2004; Padoa-Schioppa and Assad, 2006; Bouret and Richmond, 2010; Kobayashi et al., 2010; Hosokawa et al., 2013; Rudebeck, 2013; Rich and Wallis, 2016; Yun et al., 2020). Neurons in the rmCD are also known to signal incentive values of future action (Nakamura et al., 2012; Fujimoto et al., 2019). Moreover, neuronal signals in both the OFC and the rmCD have been shown to be affected by the internal states of satiety. For example, reward-specific satiety reduced OFC neuronal signals related to olfactory and gustatory stimuli (Rolls et al., 1989; Critchley and Rolls, 1996) and the subjective value for economic choice (Pastor-Bemier et al., 2021). Additionally, task-related activity of some rmCD neurons was modulated by the satiation level in the reward-size task (Fujimoto et al., 2019). However, the loss of these value- and satiety-related neuronal signals by lesions or inactivation at each stage did not affect the satiation effect on goal-directed action, as discussed above. Thus, the behavioral alterations observed in the monkeys following the OFC-rmCD disconnection in our study may have been caused by the loss of interaction of these neuronal signals through this connection.

Lesions of the OFC have been repeatedly shown to abolish normal choice adapting behavior to changes in reward value through satiety (i.e., devaluation effect)(Izquierdo et al., 2004; Machado and Bachevalier, 2007; Baxter et al., 2009; Rudebeck et al., 2013). Furthermore, inactivation studies revealed that the OFC (i.e., BA13) is essential for updating the valuation of expected reward outcomes, whereas BA11 is critical for translating this knowledge into goals (West et al., 2011; Murray et al., 2015). Several studies in rodents have also reported that satiation-induced decreases in instrumental actions were blocked by inactivation of the projection from the OFC to the dorsomedial striatum (Yin et al., 2005; Gremel and Costa, 2013; Gremel et al.,2016). Our data are consistent with these previous findings, and highlight the role of the primate OFC-rmCD pathway in the motivational adjustment of action on the basis of incentive and drive.

Inactivation of the unilateral OFC significantly shortened the reaction time and increased the total reward accumulation, which may suggest a general increase in motivation. These effects were not lateralized, as they were observed in both monkeys even though the silenced OFCs were in opposite hemispheres. Such phenomena are not similar to the deficits that we observed following OFC-rmCD disconnection—an impairment in motivational control of goal-directed action. Although the exact contribution of the OFC in one hemisphere to motivational control of behavior remains an open question, our results indicate that the specific functional connection between the OFC and the rmCD is critical for modulating behavior on the basis of the expected reward value.

In the present study, functional disconnection was attempted by expressing inhibitory DREADDs in the OFC and rmCD contralaterally, as in a crossed-lesion disconnection design (Baxter et al., 2000; Clark et al., 2013; Eldridge et al., 2016). Because the expression ofhM4Di in the rmCD mirrored that in the terminal field of OFC-rmCD projection (*cf*. Fig. 3A), the crossed chemogenetic silencing appeared to disrupt the communication between the OFC and the rmCD bilaterally. However, for anatomical disconnection, a pathway-specific chemogenetic silencing method would be ideal, such as local agonist delivery (Oyama et al., 2021) or a double viral vector system (Oguchi et al., 2021). It should also be noted here that the two DREADD actuators, CNO and DCZ, were used in our study. This was because DCZ was developed in parallel with the progress of the present experiment. Regardless of the actuator used, comparable and consistent behavioral changes specifically occurred after completion ofbilateral vector injections, confirming that this event was caused by the hM4Di-mediated functional disconnection between the OFC and the rmCD, rather than other factors such as off-target actions of metabolites.

The current findings do not exclude the possibility that the impairment observed following OFC-rmCD disconnection is caused by removing motivational processing at a third structure receiving input from OFC and rmCD. For example, the medial magnocellular part of the mediodorsal thalamus (MDmc) is known to receive input directly from the OFC (McFarland and Haber, 2002; Xiao et al., 2009) and from rmCD via the ventral pallidum (Russchen et al., 1987). Indeed, our PET data indicated the presence ofhM4Di-positive terminals in the MDmc (Fig. 3B). Furthermore, lesions ofthe MDmc have been reported to disrupt reinforcer devaluation effects in monkeys (Mitchell et al., 2007). Thus, further studies are needed to identify the pathways that contribute to OFC and rmCD communication in goal-directed behavior.

Our findings further have important implications for understanding neuropsychiatric disorders, whose symptoms can be associated with abnormal motivational control of behavior. For example, patients with OCD show a general impairment in the ability to flexibly adjust their behaviors to changes in outcome values, resulting in an over-reliance on habits (Gillan et al., 2011; Gillan and Robbins, 2014). Deficits in goal-directed control are also observed in other conditions, including major depressive disorders, substance abuse, and binge-eating disorders (Griffiths et al., 2014; Jahanshahi et al., 2015; Fettes et al., 2017). Neuroimaging studies have suggested dysfunction of the prefronto-striatal network in the pathophysiology of these disorders (Figee et al., 2013; Abe et al., 2015; Foerde et al., 2015; Voon et al., 2015; Ng et al., 2019; Sha et al., 2020). In this context, the present study provides valuable causal information about a specific neural network in primates. Future chemogenetic studies combined with functional MRI will provide a unique opportunity to link behavioral and network changes in monkeys (e.g., Hirabayashi et al., 2021), and directly compare these changes with those seen in human psychiatric disorders. As a component of this future research, it would be useful to assess obsessive-compulsive or other behaviors that are not addressed in the current study.

In summary, chemogenetic disconnection of communication between the OFC and the rmCD produced a significant impairment in the normal estimate of reward value along with satiation. Our observations are in accordance with previous evidence for neuronal signaling related to incentive and drive that has been identified in these structures during goal-directed behavior. Additionally, previous lesion and inactivation studies of these brain areas suggest a causal role of value signals related to the OFC and the rmCD in behavioral adjustment. Our results extend these previous findings and directly demonstrate that the functional connection between the OFC and the rmCD is critical for generating motivational value based on the integration of external stimuli with internal drive in monkeys. The present data also have clinical implications that could be useful for advancing our understanding of the pathophysiology of certain psychiatric disorders.

## Ackuowledgrnents

This study was supported by MEXT/JSPS KAKENHl Grant Numbers JP21K07268 (to KO), JPI5H05917, JPI5K21742, and JP20H05955 (to TM), by AMED Grant Numbers JPI8dm0307007 (to TH), JPI9dm0207077 (to MT), and JPI6dmOl07146 (to TM), by the Intramural Research Program ofNlMH/NlH/DHHS (Annual Report: ZIAMH0026 19), by the cooperative research program (2021-A-14) at the PRI, Kyoto Univ., and by National BioResource Project: “Japanese Monkeys” of MEXT, Japan. We thank J. Kamei, R. Yamaguchi, Y Matsuda, Y Sugii, T. Okauchi, T. Kokufuta for their technical assistance.

